# Analyzing the dynamics in defense/counter-defense games among hosts and pathogens

**DOI:** 10.64898/2026.05.27.728168

**Authors:** Shalu Dwivedi, Leonardo Oña, Stefan Schuster

## Abstract

In the dynamic interplay between hosts and pathogens, hosts may produce a defense compound that acts as a toxin to deter pathogen attack. Conversely, pathogens may evolve to produce a counter-defense enzyme, neutralizing the host’s toxin. This evolutionary arms race incurs costs for both parties, prompting adaptations and strategic shifts. We conceptualize this interaction as an asymmetric game, with hosts and pathogens as players, and their potential responses - defense, counter-defense, or inaction - as their strategic options. In this scenario, if the pathogen’s counter-defense enzyme is entirely effective, then the host’s toxin is rendered obsolete. However, should the host cease toxin production, the pathogen’s enzyme becomes redundant, ironically reinstating the toxin’s utility. This interaction leads to potential red-queen cycles in defense and counter-defense strategies under certain conditions, or a balanced, optimal production of toxin and enzymes by hosts and parasites, respectively. To explore this, we introduce a game-theoretical model incorporating replicator dynamics to examine temporal shifts in strategy from active (counter-)defense to non-(counter-)defense and back. In addition, we analyze compromise strategies and interpret them as bet-hedging-like. We provide a deterministic illustration of how partial defense and counter-defense generate a fitness-buffering structure in unpredictable environments and increase the geometric mean fitness of the population. In conclusion, our analysis supports the notion of continuous periodic adjustments in strategies, notably in the levels of defensive and counter-defensive measures.

## 1 Introduction

### 1.1 Background and aim of the study

Many organisms are susceptible to pathogen attack. However, the susceptibility of the host varies greatly in terms of its ability to resist or defend against the pathogen. A primary defense mechanism in hosts involves the production of chemical compounds that inhibit pathogens, such as reactive oxygen species. In response, pathogens can produce counter-defense compounds, such as superoxide dismutases, to circumvent or detoxify the host’s defense chemicals (Balloy and Chignard, 2009; Frohner *et al*., 2009; Gow *et al*., 2012; Dühring *et al*., 2015; Mackel and Steele, 2019; Ewald *et al*., 2020; Dwivedi *et al*., 2025). Further examples will be given in the next subsection. A schematic diagram of the interactions among these populations is given in Figure 1.

**Figure 1.**
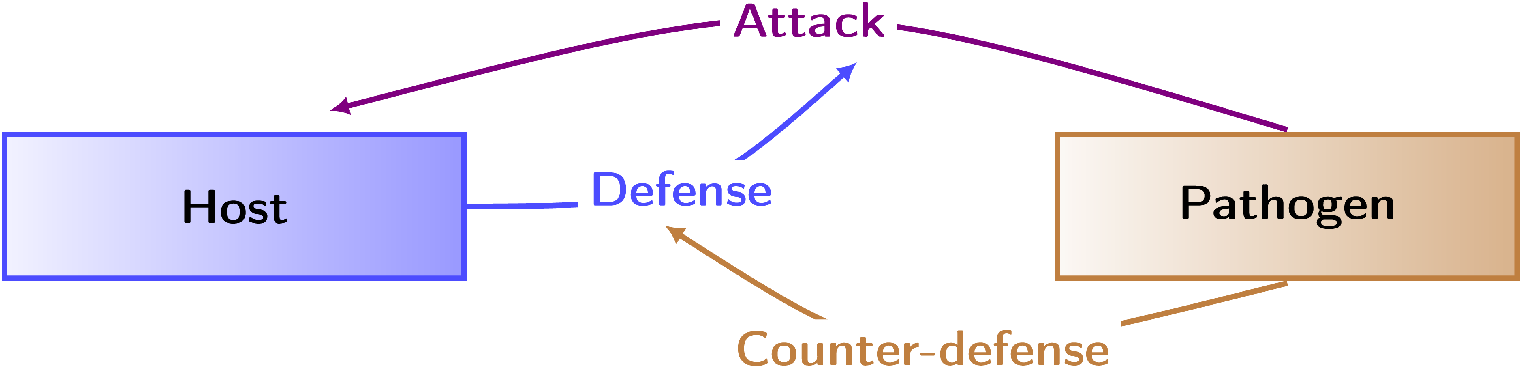
Schematic diagram of the host-pathogen interactions

**Figure 2.**
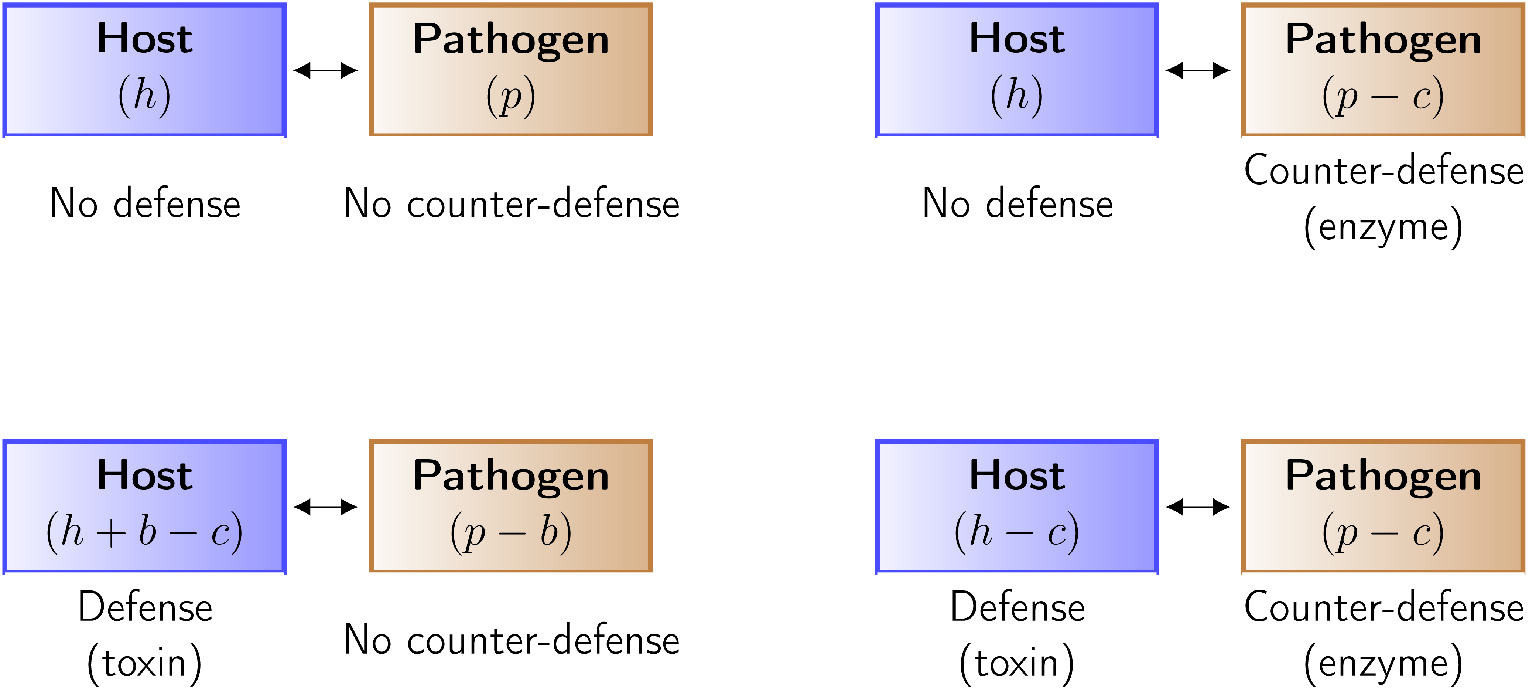
Schematic diagram of host-pathogen interactions with fitness values. The four panels correspond to the four cells in Table 1.

To better understand the interactions between hosts and pathogens, mathematical modeling and computer simulation are very helpful (Bauer *et al*., 2009; Ewald *et al*., 2020). A useful methodology is evolutionary game theory (Renaud and de Meeûs, 1991; McNickle and Dybzinski, 2013; Hummert *et al*., 2010; Hummert *et al*., 2014; Dühring *et al*., 2015; Dühring *et al*., 2017; Wu and Ross, 2016; Javarone, 2018). A game mainly contains three components: Players, strategies and payoffs. Players take part in a game, strategies are actions played by the players and payoffs are the outcome of those actions (Gintis, 2009; Smith, 1982; Tadelis, 2013). While classical game theory only deals with the final equilibrium states, it is often of interest to analyse the time behavior in reaching those states. A well-suited technique for doing so is called replicator dynamics and is based on ordinary differential equations (Smith, 1982; Weibull, 1995; Hofbauer and Sigmund, 1998; Gintis, 2009; Cressman and Tao, 2014). Such equations often lead to oscillatory solutions both for intraspecific interactions between subpopulations and for interspecific interactions (Dubey *et al*., 2024). A striking example is provided by the oscillating frequencies of different throat color (orange, blue and yellow) phenotypes in a population of side-blotched lizards, which can be modelled as a rock-paper-scissors game (Sinervo and Lively, 1996; Gintis, 2009).

Earlier, a game between defensive and non-defensive plant populations was analysed. That pairwise interaction shows coexistence between them when defended plants suffer less from herbivory pressure than non-defended plants (Augner *et al*., 1991). Later, the evolution of plant diversity including chemodiversity has been studied with the help of verbal and quantitative models, together with mathematical models and computer simulations (Chakraborty *et al*., 2023a; Thon *et al*., 2024; Kessler and Halitschke, 2009). These models also describe the interaction of plants with pollinators and herbivores. Multi-partner conflicts have been analysed in ant–plant–pollinator systems, where defensive behavior by ants reduces plant reproduction by deterring pollinators and produces parameter thresholds separating coexistence from pollinator loss (Oña and Lachmann, 2011).

Earlier, we presented a game-theoretical dynamics approach to describing such interactions, in view of the fact that the outcome for each interacting organism also depends on the action of the counterpart (Dwivedi *et al*., 2025). We used classical concepts of Game Theory, notably payoff matrix and Nash equilibrium (named after John Nash) (Gintis, 2009; Tadelis, 2013). In game-theoretical terms, the interaction under study can be considered as a generalized matching-pennies game, which has no pure Nash equilibrium (Tadelis, 2013). When counter-defense is very effective and degrades the toxin completely, then producing more toxin is not useful for the host and the latter can stop producing it. Now counter-defense is useless in absence of defense and can be stopped by the pathogen. Now, again the host can start producing toxin and defend against the pathogen (Ewald *et al*., 2020; Dwivedi *et al*., 2025).

The endless cycle of strategy switches is reflected by a mixed Nash equilibrium, implying that strategies are adopted with certain probabilities. This can be interpreted in different ways. Individual organisms may switch between strategies (red-queen cycles) or stick to one strategy, while the population subdivides into two groups showing different strategies (polymorphism). Another possible scenario consistent with a mixed Nash equilibrium is that both host and pathogen produce an intermediate amount of the toxin or enzyme (monomorphism). All three cases have also been predicted by methods based on differential equations (Buckingham and Ashby, 2022).

Here, we extend the game-theoretical approach by using replicator equations (cf. (Hofbauer and Sigmund, 1998), which allow us to analyze the temporal behavior. In particular, our goal is to figure out under which conditions oscillations occur or the two counterparts “agree” on stationary compromise levels of toxin and enzyme. We consider a game where the host and pathogen are players and choose corresponding strategies to maximize their fitness in terms of payoffs. Here, the strategies for the host are defending itself or not from the pathogen’s attack, while the strategies for the pathogen are interfering or not with the host’s defense.

We begin with the payoff matrix of the host-pathogen interaction given in Dwivedi *et al*. (2025) and as an extension, we establish a game-theoretical model with replicator equations, in which the variables are defined as the relative frequencies of strategies (Smith, 1982; Weibull, 1995; Hofbauer and Sigmund, 1998; Gintis, 2009; Cressman and Tao, 2014).

An alternative approach in game theory is based on population dynamics (Hofbauer and Sigmund, 1998; Neumann and Schuster, 2007; Křivan *et al*. 2018). In Subsection 2.2, we present the defense/counter-defense game with the help of Lotka-Volterra equations, including the population densities as variables. Both types of equations represent systems of non-linear first-order differential equations to describe the dynamics of the game in the continuous case (Taylor and Jonker, 1978). By determining the stability and asymptotic behavior of the game (Murray, 2002; Otto and Day, 2007; Ogata, 2010; Chakraborty *et al*., 2024) in these two representations, we will figure out whether self-sustained oscillations between the above-mentioned strategies occur.

Previously, we found partial defense and counter-defense to be stable steady-state strategies for appropriate parameter values of the dose-response curve (Dwivedi *et al*., 2025). These intermediate actions can be interpreted as bet-hedging strategies. This can be explained as follows (Cohen *et al*., 1966; Shemesh *et al*., 2013). Investing the full capacity in defense reduces growth too much if no pathogen attacks, while Investing nothing in defense implies serious harm or even death if a massive attack happens. Thus, it may be favorable to maintain a permanent, partial investment in defense or analogously into counter-defense. In Subsection 2.3, we analyze the strategies of the host and pathogen in view of bet-hedging. This is an adaptive strategy in which an individual’s fitness decreases but the fitness of the total population is maximized in an uncertain environment (Seger and Brockmann, 1987; Olofsson *et al*., 2009; Starrfelt and Kokko, 2012; Botero *et al*., 2014; Grimbergen *et al*., 2015). In particular, a risk-spreading strategy, providing a long-term fitness gain in unpredictable environments, is considered as a bet-hedging strategy (de Jong *et al*., 2011). An illustrative example of bet-hedging is seed dormancy, where the germination of seeds in desert plants varies within the population. While some seeds germinate, others can remain dormant for years. If a harsh weather condition (such as drought) kills the germinated seedlings, the dormant seeds ensure that the lineage continues by germinating in the future under favorable weather conditions (El-Keblawy and Gairola, 2017).

### 1.2 Examples of defense and counter-defense

To provide more biological background, we now mention several examples of defense and counter-defense. A multitude of such interactions have been studied in nature, for example, the interplay between macrophages and *Candida albicans* (Gow *et al*., 2012; Dühring *et al*., 2017), between plants and herbivores (Textor and Gershenzon, 2009; Jeschke *et al*., 2017; Chakraborty *et al*., 2023b), and between bacteria and fungi (Hogan and Kolter, 2002); Stöckli *et al*., 2017). For instance, defensins are antimicrobial peptides (AMP) against pathogenic microbes, such as viruses, bacteria and fungi (Gao *et al*., 2021). One example of a defensin is the human AMP LL37 (Burton *et al*., 2009). Many bacteria possess counter-defense mechanisms, such as the outer membrane protease omptin from *Escherichia coli*, which cleaves LL37, leading to loss of bactericidal activity (Thomassin *et al*., 2012). Plants of the Brassicaceae family defend against insect herbivores by producing toxic isothiocyanates (ITCs) through hydrolysis of glucosinolates (GLSs) (Textor and Gershenzon, 2009; Jeschke *et al*., 2017; Chakraborty *et al*., 2023b). However, to avoid the toxicity of ITCs, some specialist insect species evolved preemptive counter-defense techniques, such as breaking down GLSs before hydrolysis or redirecting the hydrolysis process to produce less toxic products or sequestering GLS before hydrolysis (Jeschke *et al*., 2017; Chakraborty *et al*., 2023b). For example, the diamondback moth (*Plutella xylostella*) desulfates GLSs before hydrolysis (Ratzka *et al*., 2002); the large cabbage white moth (*Pieris rapae*) redirects GLS hydrolysis by nitrile-specifier protein (NSP), which leads to the formation of less toxic nitriles (Wittstock *et al*., 2004); the horseradish flea beetle (*Phyllotreta armoraciae*) sequesters GLSs in its gut before hydrolysis (Sporer *et al*., 2021). Similarly, white mold (*Sclerotinia sclerotiorum*) fungi detoxify ITCs into harmless amines by using ITC hydrolase enzymes (Chen *et al*., 2020).

In a bacterial-fungal interaction, *Bacillus thuringiensis* defends against the coprophilous fungus *Coprinopsis cinerea* (Stöckli *et al*., 2017) and *Pseudomonas aeruginosa* defends against *C. albicans* (Hogan and Kolter, 2002) in the struggle for nutrients by producing biofilm and antifungal molecules. They use coordination by the quorum sensing system, and interference with this bacterial communication system is used as a counter-strategy by the fungus in this interaction (Stöckli *et al*., 2017).

The host defense pathways show cooperation, complementation, and compensation strategies against pathogenic bacteria and viruses (Nish and Medzhitov, 2011). Innate immune defense against the different phenotypes of *Aspergillus fumigatus* fungi is characterized by recognition, phagocytosis, intracellular and extracellular killing (Balloy and Chignard, 2009). The intact lung epithelium prevents the adhesion of *A. fumigatus conidia* (Mackel and Steele, 2019). Another study describes the strategies used by phages (in the family *Siphoviridae*) to overcome resistance mechanisms of the bacterial host (*Escherichia coli*), including inhibition of adsorption and restriction–modification in phage–host interactions (Samson *et al*., 2013). An experimental result shows that the T4 phage (subfamily *Tevenvirinae* of the *Straboviridae*) can evolve to overcome a phage-defensive toxin-antitoxin system (toxIN) in *Escherichia coli* (Srikant *et al*., 2022).

Another example of host-bacterial interactions is the innate immune system’s production of reactive nitrogen species as a defense against pathogens. Pathogenic bacteria such as *Mycobacterium tuberculosis* (Bhat *et al*., 2017) and *Salmonella* (Zhao *et al*., 2017) slow down the expression of NOS via RNA interference and binding of PPE2 to the TATA box of the iNOS gene (Bhat *et al*., 2017; Zhao *et al*., 2017). Another bacterium, *Francisella tularensis*, inhibits iNOS expression by suppressing IP-10 production and IFN*γ* induced STAT-1 signaling (Parsa *et al*., 2008). An example of conflict between endosymbionts and their hosts was discussed where the mammalian immune system interacts with parasites *Wolbachia* including cytoplasmic incompatibility (CI) infection (Hammerstein *et al*., 2005).

Oscillatory changes in strategies have been analyzed previously. Examples of such a behavior in intraspecific interactions are provided by the oscillating frequencies of different phenotypes of lizards Sinervo *et al*., 2000) (see above) and of two cichlid fish’s phenotypes in lake Tanganyika, where the abundance of individuals with left- or right-handedness of mouth opening depends on frequency-dependent natural selection (Hori *et al*., 1993). As for predator-prey interactions, it is well known that oscillations are the typical behavior (Yoshida *et al*., 2003). Moreover, also host-parasite interactions can give rise to such a dynamics (Dybdahl and Lively, 1998; Buckingham and Ashby, 2022).

## 2 Methodology and Results

We consider a game where the host and the pathogen play their actions as players. Host has two strategies, namely: No defense (ND) and Defense (D) (see Augner *et al*., 1991, for the case of plants). Similarly, pathogen has two strategies, namely: No counter-defense (NCD) and Counter-defense (CD) (Dwivedi *et al*., 2025).

Table 1 gives the payoffs (fitness values) for host and pathogen.

**Table 1.** Fitness matrix for host and pathogen (Dwivedi *et al*., 2025).

| Host, Pathogen | No counter-defense (NCD) | Counter-defense (CD) |
| --- | --- | --- |
| No defense (ND) | $(h, p)$ | $(h, p - c)$ |
| Defense (D) | $(h + b - c, p - b)$ | $(h - c, p - c)$ |

Here, we denote the payoff values of the host and pathogen without any defense and counter-defense as (*h, p*), the cost of defense or counter-defense as *c* and the benefit as *b* (Table 1) (Dwivedi *et al*., 2025). The game is an asymmetric two-player two-strategy game.

The Nash equilibrium can simply be determined from Table 1 by iteration, until a set of strategies is reached where each player using it obtains a locally maximum payoff, given the other player’s strategy (Dwivedi *et al*., 2025). We distinguish two cases, which differ in the order relations between the payoffs. For the high-benefit case (*b* > *c*), both the host and the pathogen have an incentive to switch their strategies in every cell of the payoff matrix, so no pure Nash equilibrium occurs here.

In contrast, for the low-benefit case (*b* < *c*), they always receive less payoff if they produce defense or counter-defense as the cost is higher than the benefit. So, the Nash equilibrium is ‘ND/NCD’ (Table 1). They do not have any incentive to change these strategies (Dwivedi *et al*., 2025).

Maynard Smith and Price adapted the concept of Nash equilibrium for biological contexts, formalizing it as the evolutionarily stable strategy (ESS) (Smith and Price, 1973). A strategy is an ESS if a whole population using that strategy cannot be invaded by a small group with a mutant genotype (Smith and Price, 1973; Gintis, 2009). As an extension, in the first model described in Subsection 2.1, we apply the method of replicator dynamics (Smith, 1982; Weibull, 1995; Hofbauer and Sigmund, 1998; Gintis, 2009; Cressman and Tao, 2014), to see how the host and the pathogen change their strategies over time. We use the fitness matrix given in Table 1 and establish Model 1 to explain switching strategies of (no)defense and (no)counter-defense in host-pathogen interactions. The second model is formed with the Lotka-Volterra equations, given in Subsection 2.2 (Hofbauer and Sigmund, 1998; Gintis, 2009; Ogata, 2010). There, we analyze the population using different strategies changes over time. Further we perform the stability analysis for both models. The third model is the benefit-cost model based on Michaelis-Menten kinetics (Simms and Rausher, 1987; Siemens *et al*., 2010), mentioned in Subsection 2.3. It can be interpreted in view of bet-hedging strategies. For the third model, we first determine the fitness of the host-pathogen population. Thereafter, we calculate the arithmetic and geometric mean fitness of host and pathogen, respective to their capability of defense and counter-defense production.

The parameter values used in the Models (1, 2, 3) are given in the Table 2.

**Table 2.**
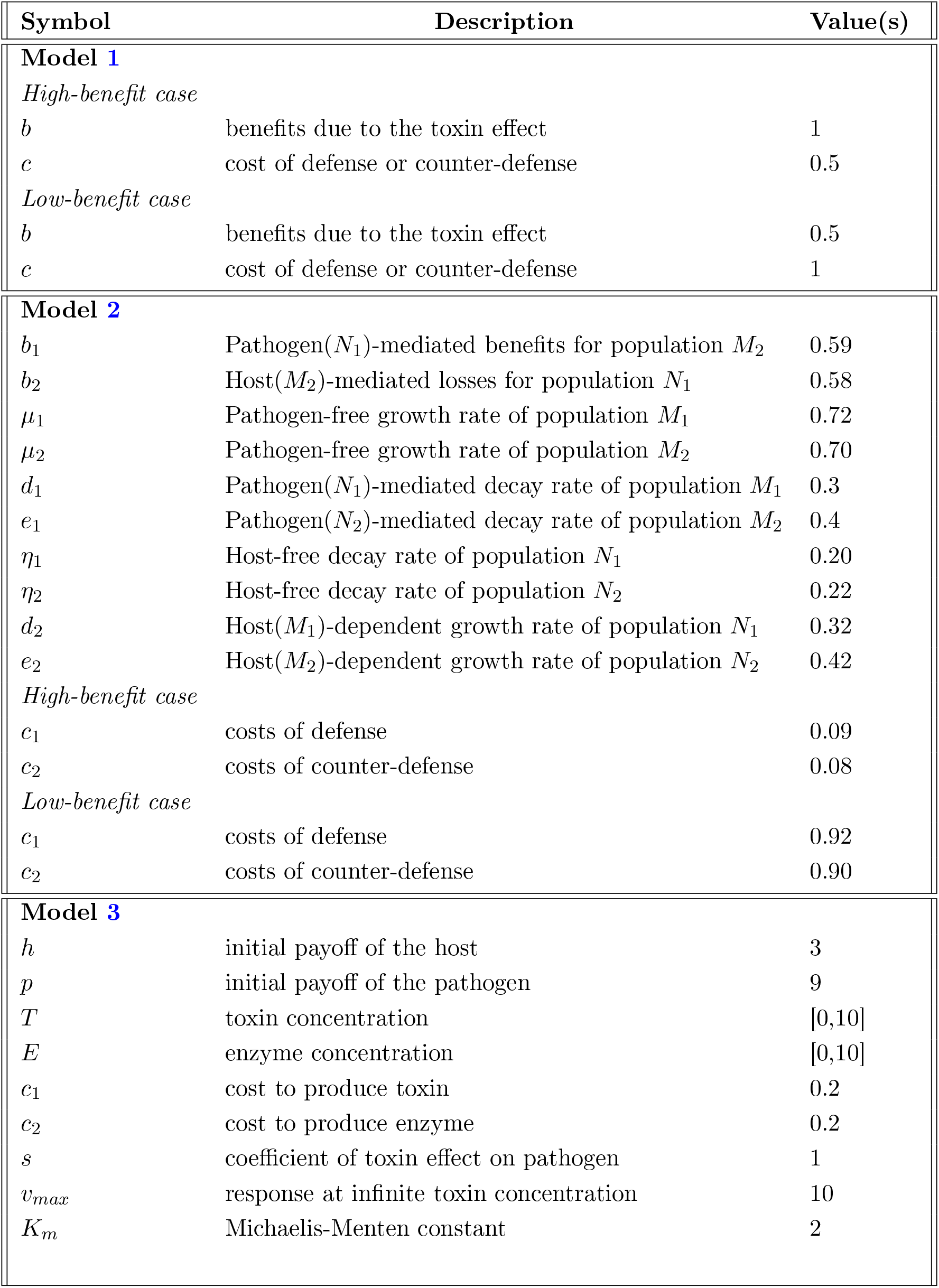
Parameter values used in Model 1, Model 2 and Model 3. (*M*_1_, *M*_2_) are hosts population that choose the no defense and defense strategy, respectively, and (*N*_1_, *N*_2_) are pathogens population that choose the no counter-defense and counter-defense strategy, respectively.

### 2.1 Interaction between individuals

Here, we consider the interaction between an individual host and an individual pathogen. Let *x* and *y* denote the frequencies of the defense (D) strategy used by the host and of the counter-defense (CD) strategy used by the pathogen, respectively. Thus, (1 − *x*) and (1 − *y*) are the frequencies of the inaction strategies.

Then, the expected payoffs read as follows. Choosing strategy ‘No defense’ (ND),

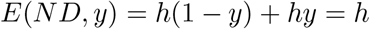

Choosing strategy ‘Defense’ (D),

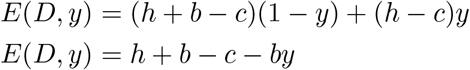

Choosing strategy ‘No counter-defense’ (NCD),

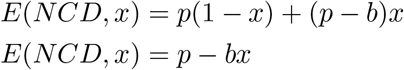

Choosing strategy ‘Counter-defense’ (CD),

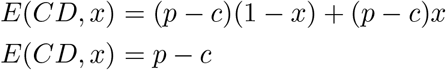

Average payoff of the host *E*(*ND, D*),

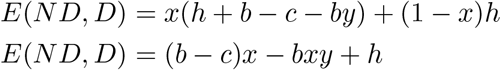

Average payoff of the pathogen *E*(*NCD, CD*),

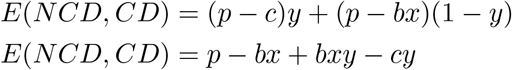

So, the replicator equations for the payoff matrix are:

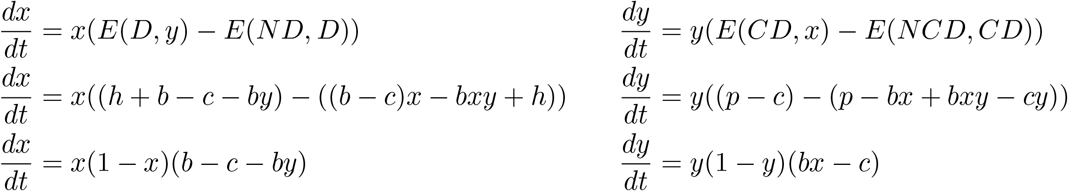

#### Model 1

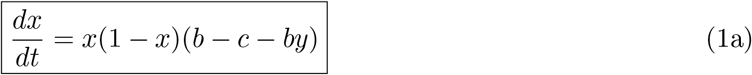

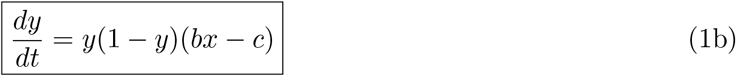

The plots of Model 1 for the high- and low-benefit case are given in Figure 3.

**Figure 3.**
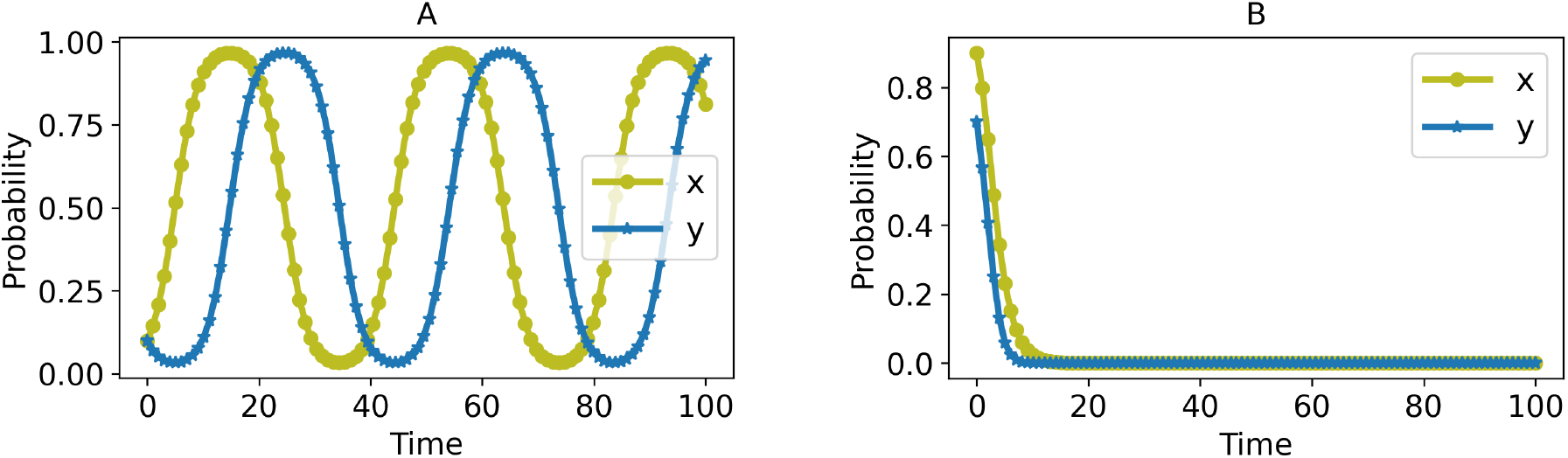
Time course of Model 1. (A) High-benefit case, parameter values: *b* = 1, *c* = 0.5. (B) Low-benefit case, parameter values: *b* = 0.5, *c* = 1. *x* and *y* are the frequencies by which an individual host and pathogen play defense and counter-defense strategies, respectively.

Here, the host and the pathogen switch their strategies continuously with some probabilities (Figure 3A) when the benefit is higher than the cost and they prefer to choose no defense and no counter-defense as the cost exceeds the benefit (Figure 3B).

The equilibrium points of Model 1 are (0, 0), (1, 0), (0, 1), (1, 1) and 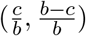, where,

- (0,0) represents ‘No defense, No counter-defense’,
- (0,1) represents ‘No defense, Counter-defense’,
- (1,0) represents ‘Defense, No counter-defense’,
- (1,1) represents ‘Defense, Counter-defense’ and
- 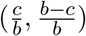 represents ‘Defense, Counter-defense’ with some probability.

The equilibria involving values of zero and/or one arise because the differential equations involve the terms *x*(1 − *x*) and *y*(1 − *y*). The equilibrium point 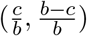 represents the mixed strategy. In other words, we can consider it as ‘Partial defense, Partial counter-defense’ strategies (Dwivedi *et al*., 2025). It is only relevant in the high-benefit case (*b* > *c*) because otherwise, it is not situated in the positive quadrant.

We can perform a stability analysis (Murray, 2002; Otto and Day, 2007; Chakraborty *et al*., 2024) of Model 1 at the equilibrium points using the Jacobian matrix of equations (1a), (1b):

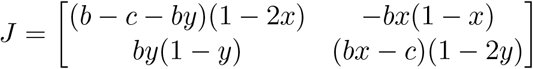

#### 2.1.1 High-benefit case (*b* > *c*)

1. At (0,0), 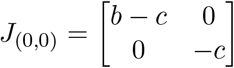 with eigenvalues (*λ*) = *−c, b* − *c*. These correspond to a saddle point, which is unstable. (Figure 4A).
2. At (0,1), 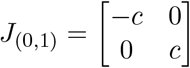 with eigenvalues (*λ*) = ±*c*, corresponding to a saddle point (Figure 4B)
3. At (1,0), 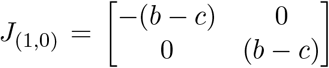 with eigenvalues (*λ*) = ±(*b* − *c*), corresponding to a saddle point (Figure 4C).
4. At (1,1), 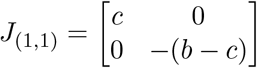 with eigenvalues (*λ*) = *c, −*(*b* − *c*), corresponding to a saddle point (Figure 4D).
5. At 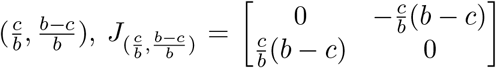 with eigenvalues 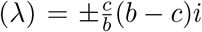, which are purely imaginary, leading to a marginally stable oscillation (Figure 6). This implies that the amplitude and period of the oscillations depend on initial conditions.

The stability analysis of the equilibrium points is shown in Figures 4-6.

**Figure 4.**
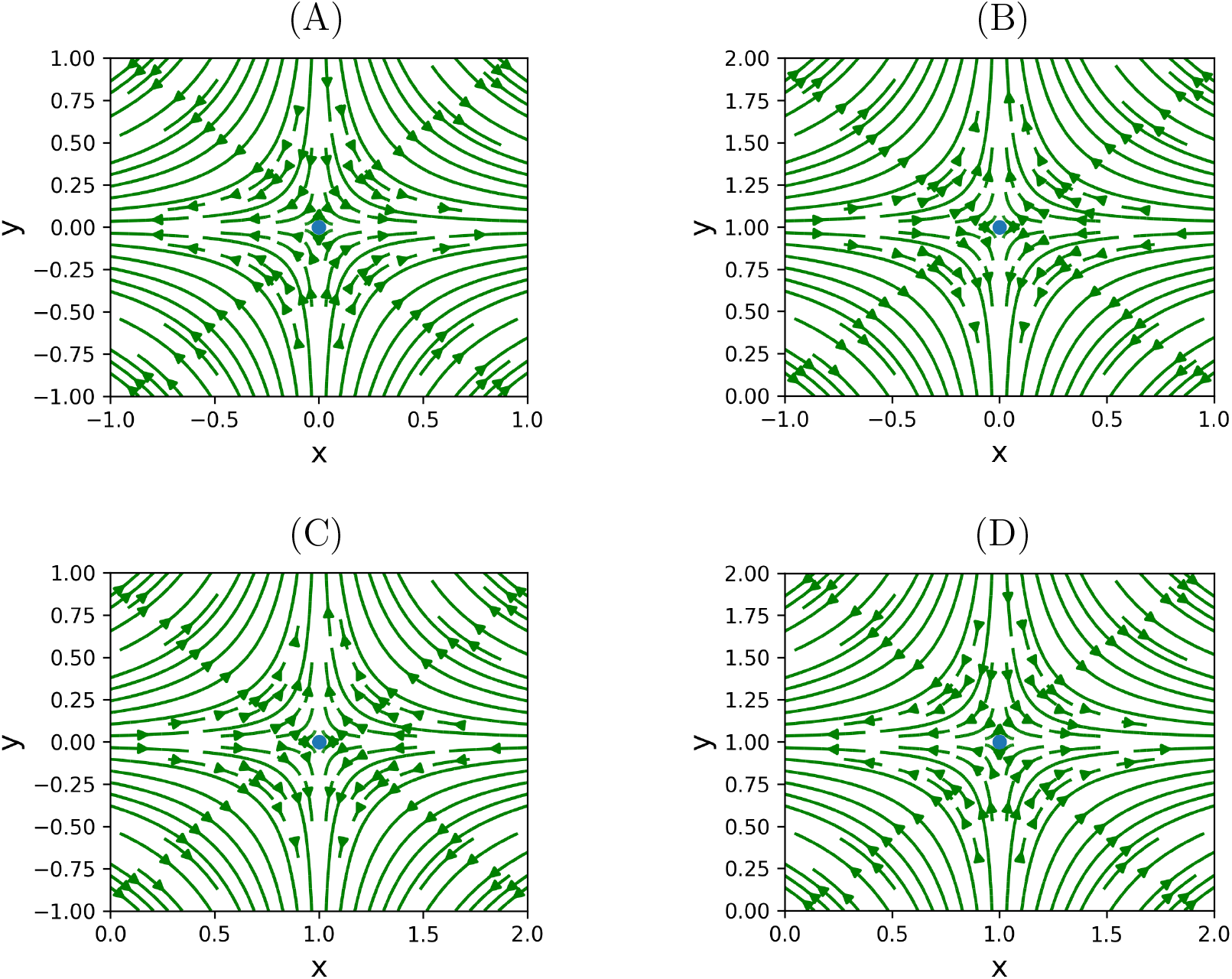
Phase portrait of equilibrium points for the high-benefit case (*b* > *c*). (A) (0,0): ‘No defense/No counter-defense’, (B) (0,1): ‘No defense/ Counter-defense’, (C) (1,0): ‘Defense/No counter-defense’ (D) (1,1): ‘Defense/Counter-defense’, saddle point

**Figure 5.**
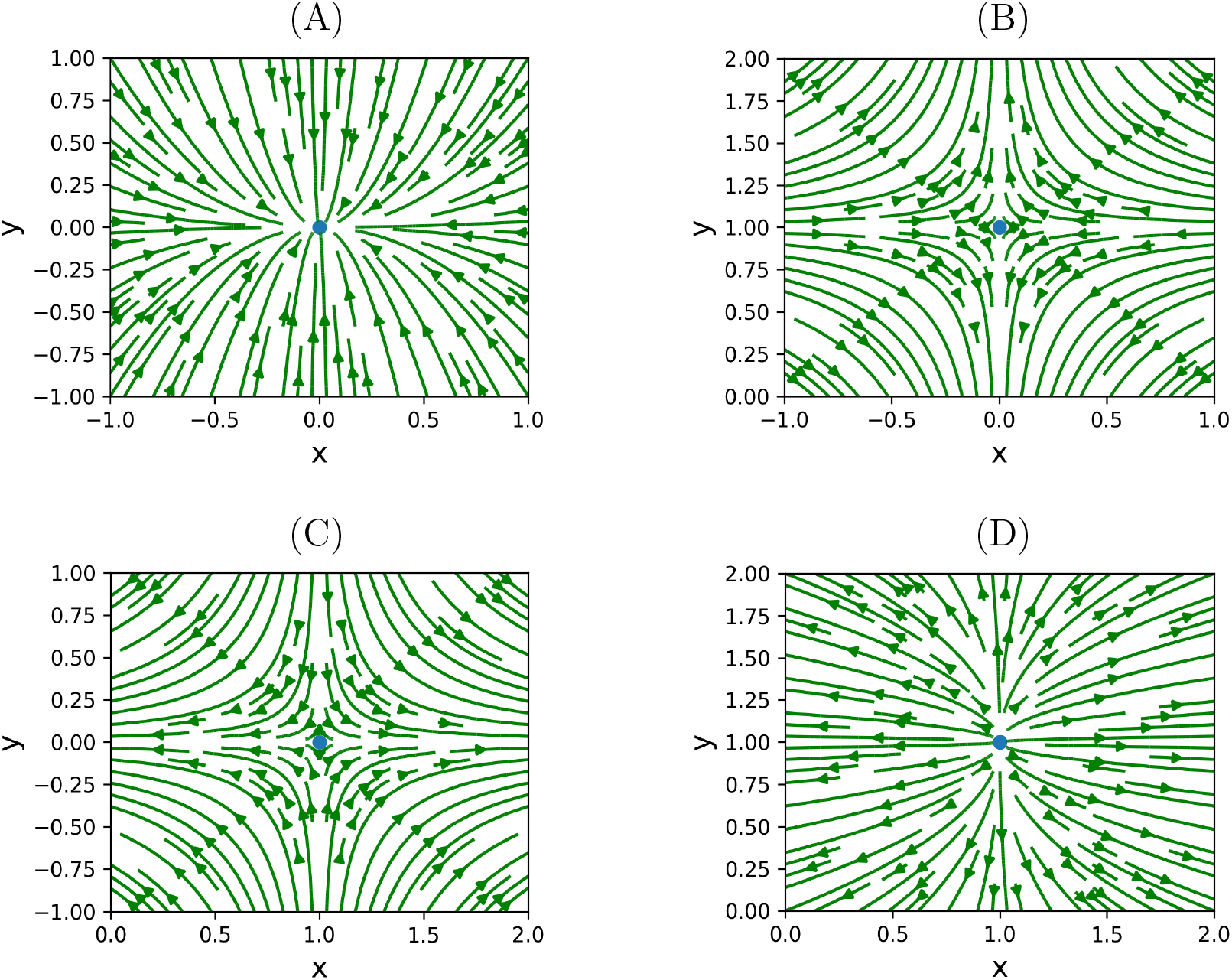
Phase portrait of equilibrium points for the low-benefit case (*b* < *c*). (A) (0,0): ‘No defense/No counter-defense’, stable node. (B) (0,1): ‘No defense/Counter-defense’, saddle node, (C) (1,0): ‘Defense/No counter-defense’, saddle node. (D) (1,1): ‘Defense/Counter-defense’, unstable node.

**Figure 6.**
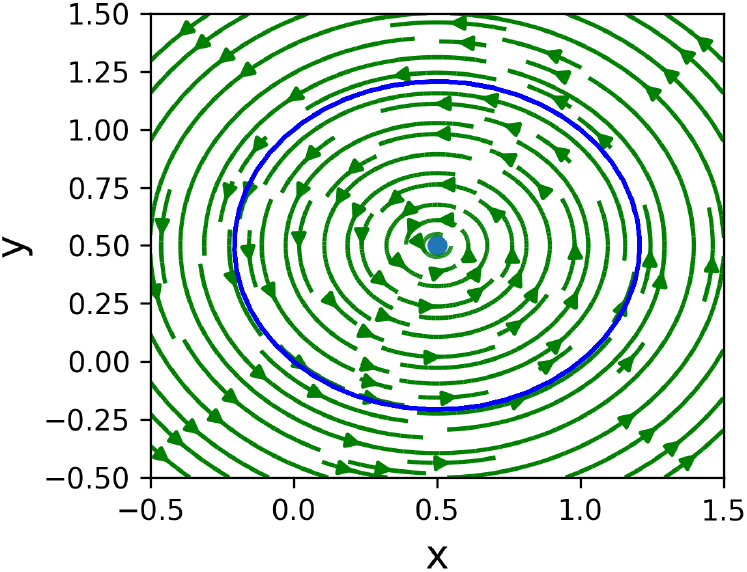
Phase portrait of equilibrium point 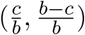 for the high-benefit case (*b* > *c*), the equilibrium point is a center.

#### 2.1.2 Low-benefit case (*b* < *c*)

1. At (0,0), 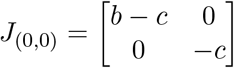 with eigenvalues (*λ*) = *−c, b* − *c*. These correspond to a stable node, which is asymptotically stable (Figure 5A).
2. At (0,1), 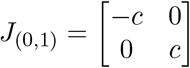 with eigenvalues (*λ*) = ±*c*. These correspond to a saddle point that is unstable (Figure 5B).
3. At (1,0),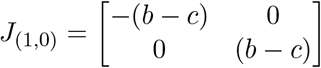 with eigenvalues (*λ*) = ±(*b* − *c*), which is a saddle point (Figure 5C).
4. At (1,1), 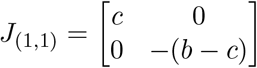, the eigenvalues (*λ*) = *c, −*(*b* − *c*), which is an unstable node (Figure 5D).

Table 3 gives a summary of the stability analysis of the equilibrium points in the individual interaction.

**Table 3.**
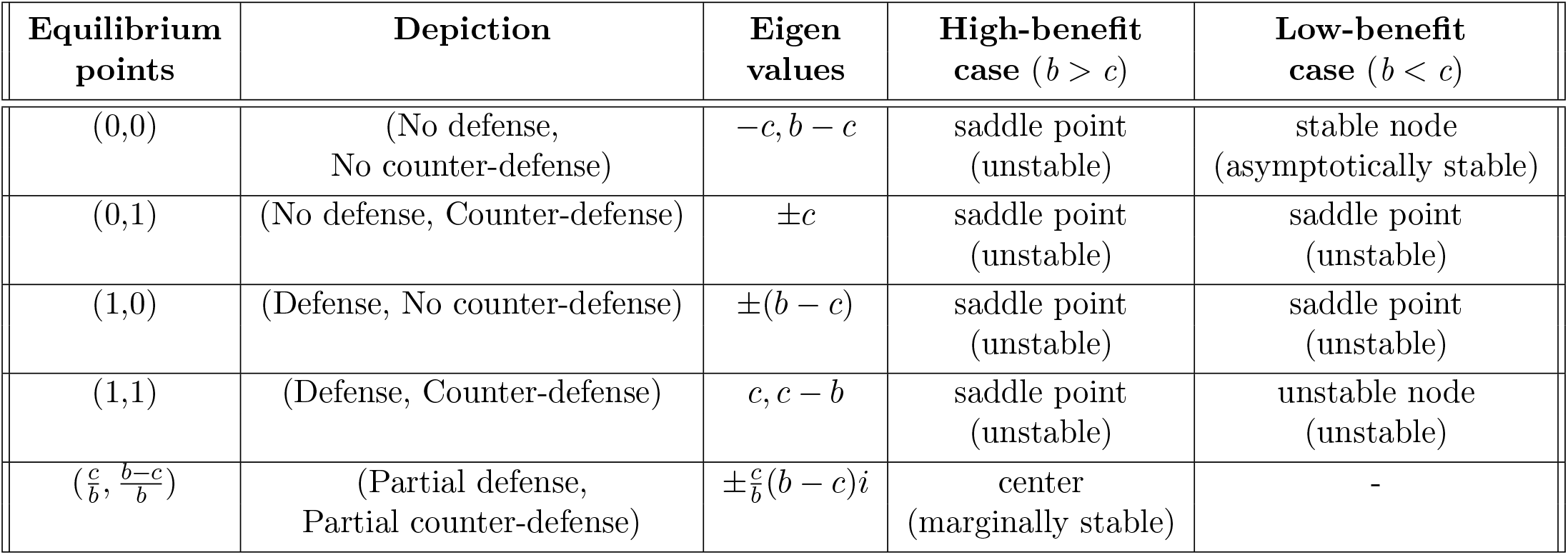
Summary of stability analysis in individual interaction. *b*: benefit, *c*: cost

From the phase portraits (Figures 4-6) and Table 3, it is clear that the ‘no defense’ and ‘no counter-defense’ strategy pair is asymptotically stable only in the low-benefit case. In contrast, the strategy pair ‘defense’ and ‘counter-defense’ with some probability is marginally stable in the high-benefit case.

### 2.2 Interactions among populations

As mentioned in the Introduction, our model 2 is based on population dynamics (Hofbauer and Sigmund, 1998; Neumann and Schuster, 2007; Křivan *et al*. 2018). Here, we consider a game among host and pathogen populations, where both defending and non-defending hosts interact with both counter-defended and non-counter-defended pathogens.

#### Model 2

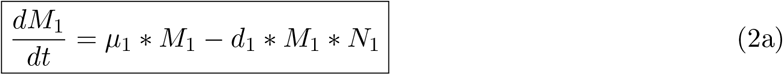

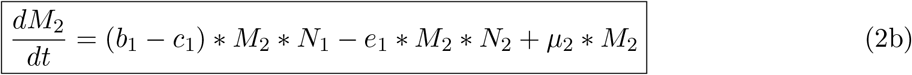

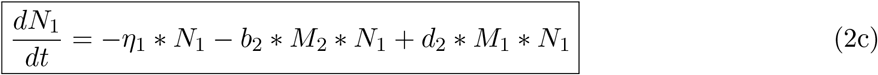

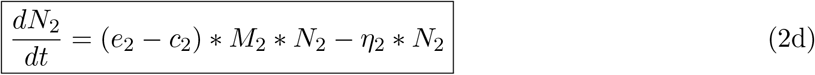

where, (*M*_1_, *M*_2_) denote the population densities of the hosts that choose the no defense and defense strategy, respectively, and (*N*_1_, *N*_2_) population densities of the pathogens that choose the no counter-defense and counter-defense strategy, respectively. (*c*_1_, *c*_2_) are the costs of defense and counter-defense for the host and pathogens, respectively. (*b*_1_, *b*_2_) are benefits for *M*_2_ and losses for *N*_1_, respectively. The populations (*M*_1_, *M*_2_) increase at rates (*µ*_1_, *µ*_2_) in the absence of pathogens and decrease with rates (*d*_1_, *e*_1_) in the interaction with (*N*_1_, *N*_2_), respectively. In contrast, the populations (*N*_1_, *N*_2_) decrease with rates (*η*_1_, *η*_2_) in the absence of hosts and increase with rates (*d*_2_, *e*_2_) in the interaction with (*M*_1_, *M*_2_), respectively.

Here, cost of defense and counter-defense are only used during the host-pathogen interaction. In the free environment, these population save the costs and reallocate resources towards unrestricted growth.

The time courses obtained by Model 2 for the high- and low-benefit cases are illustrated in Figures 7 and 8.

**Figure 7.**
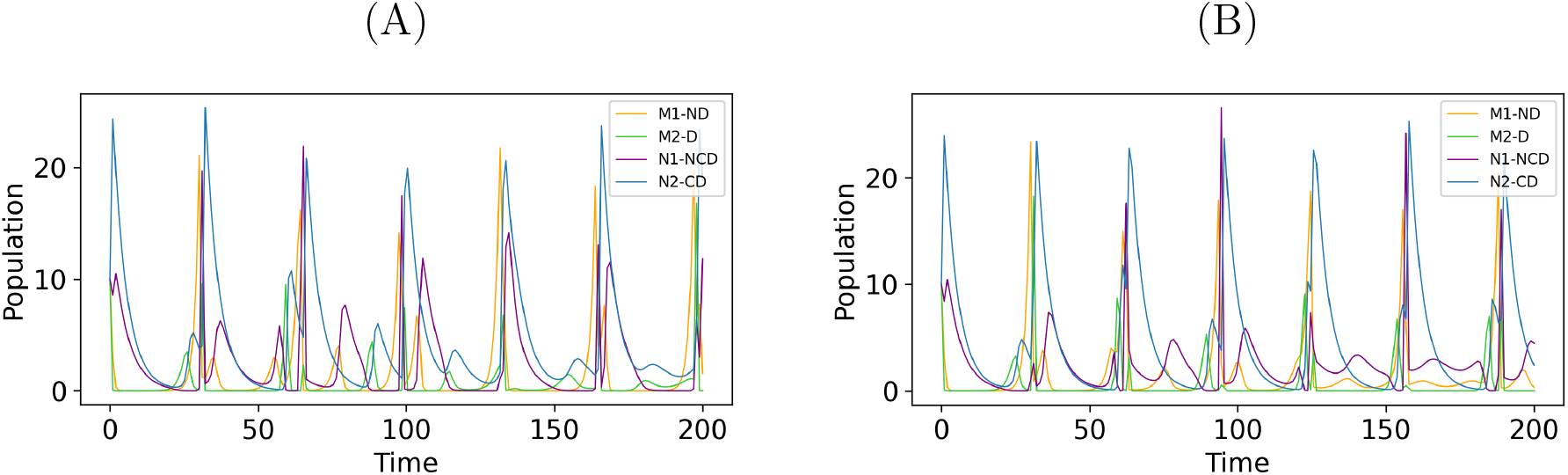
Time course of Model 2 for the high-benefit case. (A) Cost of defense and counter-defense are different, *c*_1_ = 0.09, *c*_2_ = 0.08. (B) Cost of defense and counter-defense are the same, *c*_1_ = *c*_2_ = 0.09. Parameter values are listed in Table 2.

**Figure 8.**
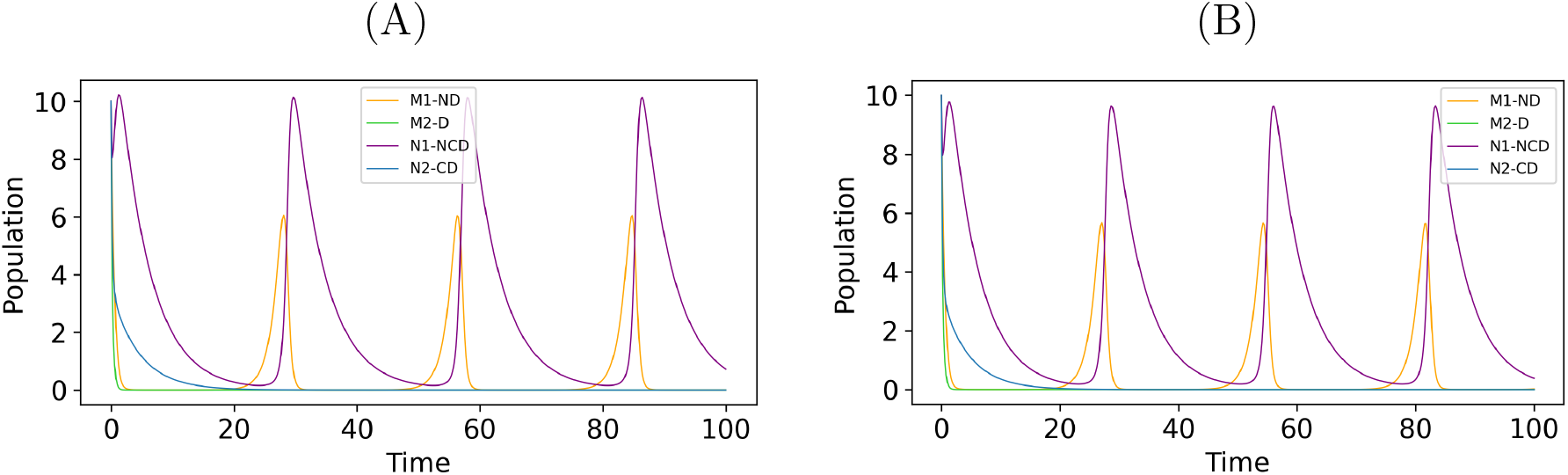
Time course of Model 2 for the low-benefit case. (A) Cost of defense and counter-defense are different, *c*_1_ = 0.92, *c*_2_ = 0.90. (B) Cost of defense and counter-defense are the same, *c*_1_ = *c*_2_ = 0.9. Parameter values are listed in Table 2.

Equilibrium points of Model 2 are 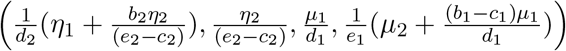 and 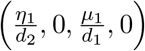. We can perform a stability analysis (Murray, 2002; Otto and Day, 2007; Ogata, 2010) of Model 2 at the equilibrium points. Since the eigenvalues are imaginary here for (*b*_1_ > *c*_2_), (*e*_2_ > *c*_2_), 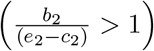 and 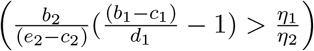, the equilibrium point 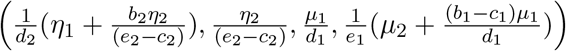 is center i.e. marginally stable (Ogata, 2010). 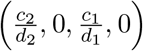 is stable when (*b*_1_ < *c*_2_) and 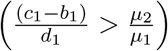.

### 2.3 Host’s and Pathogen’s responses as bet-hedging-like strategies

Fitness generally corresponds to reproductive fitness (Wild and Taylor, 2004). We subdivide the offspring of hosts and pathogens into three categories with different characteristics in the context of producing a defense chemical (e.g., toxin). We refer to these categories as phenotypes and determine their fitness by their benefit due to production of the toxin. We divide these phenotypes into conservative “bet-hedgers” (mixture of defense strategies), pure defense and no defense subpopulations. The bet-hedger populations of the host and the pathogen use defense or counter-defense with varying amounts of toxin or enzyme, respectively. Bet-hedging is a strategy in which an individual’s fitness decreases in a fluctuating environment to maximize the population’s long-term geometric mean fitness across generations (Seger and Brockmann, 1987; Olofsson *et al*., 2009; Starrfelt and Kokko, 2012; Botero *et al*., 2014; Grimbergen *et al*., 2015; Haaland *et al*., 2019). Compared to other strategies, a bet-hedging strategy is also defined as a risk-spreading strategy based on phenotypic heterogeneity that provides a long-term fitness advantage in an uncertain time-varying environment (de Jong *et al*., 2011;Shemesh *et al*., 2013). We consider the interaction with the pathogens as the environment for the hosts. That environment consists of pathogens that produce or do not produce enzymes as a counter-defense against hosts. Here, we consider three kinds of phenotype in hosts:

- Phenotype 1, non-defensive (ND) against pathogens,
- Phenotype 2, uses bet-hedging (BH) strategy, and
- Phenotype 3, always produces defense (D).

Similarly, three kinds of phenotype in pathogens:

- Phenotype 1, non-counter-defensive (NCD) against hosts,
- Phenotype 2, uses bet-hedging (BH) strategy, and
- Phenotype 3, always produces counter-defense (CD).

Phenotype 1 produces no defense or counter-defense at all and saves the costs of toxin or enzyme production. In contrast, Phenotype 3 invests a cost for toxin or enzyme production. In contrast, Phenotype 2 uses mixed strategies, which can be interpreted as bet-hedging strategies. That is, it uses a partial amount of defense chemical when required against an uncertain environment. It reduces the cost of production while retaining partial protection. It does not use toxin completely at a time.

Here we determine the fitness values for the host and pathogen using Michaelis-Menten kinetics (Simms and Rausher, 1987; Siemens *et al*., 2010).

#### Model 3

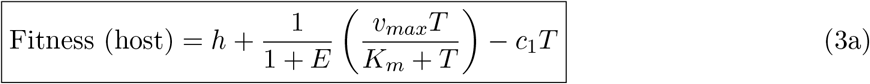

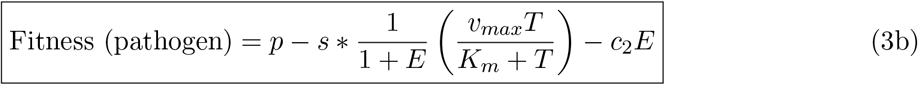

where *T*, toxin concentration; *E*, enzyme concentration; (*c*_1_, *c*_2_), cost to produce toxin and enzyme; *s*, coefficient of toxin effect on pathogen; *v*_*max*_, response at infinite toxin concentration; *K*_*m*_, Michaelis-Menten constant. The fitness values of the host and pathogen that never produce any defense and counter-defense chemicals are *h* and *p*, respectively.

Model 3 is influenced by a dose-response model (Gadagkar and Call, 2015; Dwivedi *et al*., 2025) and the benefit-cost model (Simms and Rausher, 1987; Siemens *et al*., 2010) proposed earlier. It is a graphical model that includes a Michaelis–Menten curve and linear costs.

Figure 9 provides the fitness of all three phenotypes of the host (Figure 9A) and all three phenotypes of the pathogen (Figure 9B). These values are based on the concentrations of enzyme and toxin, respectively. In the absence of enzyme, the fitness of phenotype 2 (BH) and phenotype 3 is maximum and it starts decreasing as the enzyme concentration increases (Figure 9A). In Figure 9B, it can be seen that phenotype 2 (BH) has the largest fitness in comparison to the other phenotypes unless the toxin concentration is very low.

**Figure 9.**
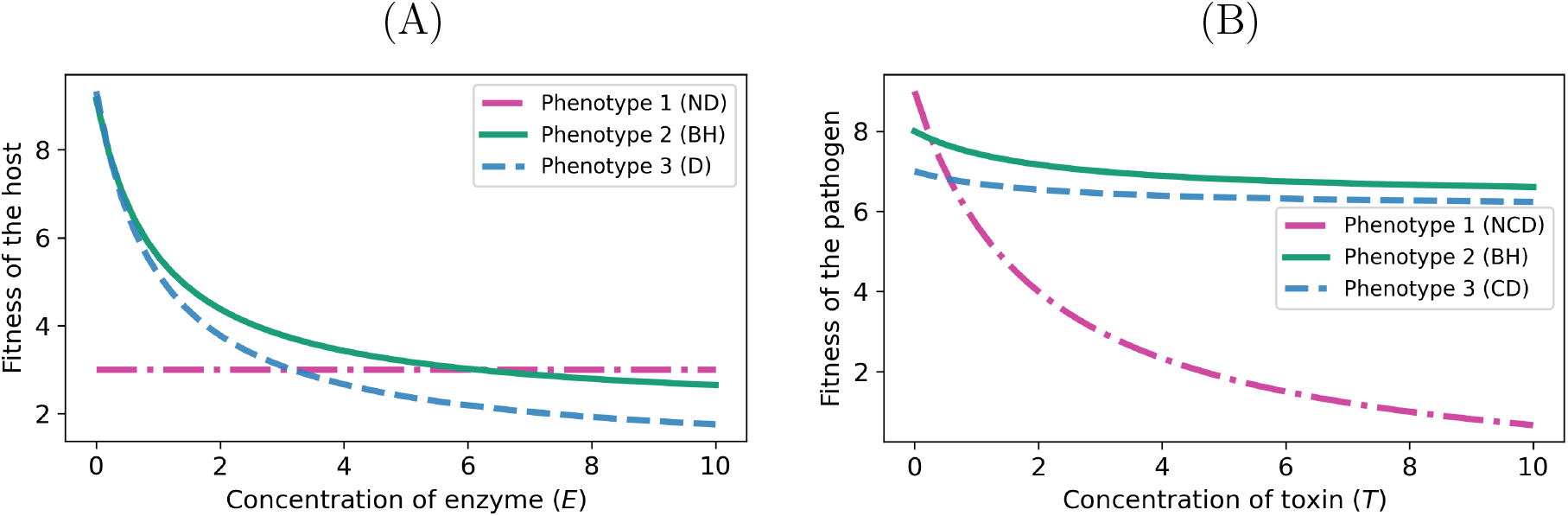
Fitness values of the host and pathogen depending on enzyme and toxin, respectively. (A) Host fitness. *T* = 0 (Phenotype 1), *T* = 5 (Phenotype 2), *T* = 10 (Phenotype 3) (B) Pathogen fitness *E* = 0 (Phenotype 1), *E* = 5 (Phenotype 2), *E* = 10 (Phenotype 3). ND: no defense, D: defense, BH: bet-hedging, NCD: no counter-defense, CD: counter-defense. Parameter values: *h* = 3, *p* = 9, *v*_*max*_ = 10, *K*_*m*_ = 2, *c*_1_ = *c*_2_ = 0.2, *s* = 1

Tables 4 and 5 give fitness values for the host and pathogen for Model 3. The values are determined from Figure 9. We assume that the initial fitness of host and pathogen is (6, 6) without any interaction. When a pathogen attacks the host, it exploits the host and earns a gain. Now the fitness value for the host is 3 (Table 4) and for the pathogen is 9 (Table 5). We simulate the fitness of all three phenotypes of the host (eq. 3a) and pathogen (eq. 3b) of Model 3. The values in the fitness matrix (Tables 4 and 5) are determined from Figures 9A and 9B, respectively. All parameters used here are arbitrary. The individual fitness of the host’s phenotype 3 is higher in interaction with the pathogen’s phenotype 1 (Table 4). Similarly, the individual fitness of the pathogen’s phenotype 1 is higher in interaction with the host’s phenotype 1 (Table 5).

**Table 4.** Fitness matrix for host population from Figure 9A. ND: no defense, D: defense, BH: bet-hedging.

| Host | Pathogen |  |  |
| --- | --- | --- | --- |
|  | Phenotype 1 (NCD) | Phenotype 2 (BH) | Phenotype 3 (CD) |
| Phenotype 1 (ND) | 3 | 3 | 3 |
| Phenotype 2 (BH) | 9.14 | 3.19 | 2.65 |
| Phenotype 3 (D) | 9.33 | 2.39 | 1.76 |

**Table 5.**
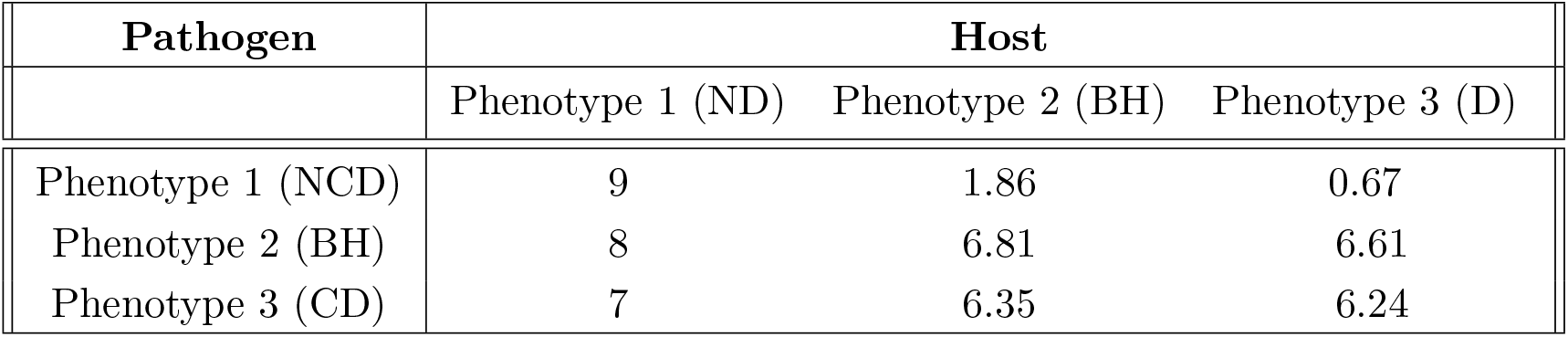
Fitness matrix for pathogen population from Figure 9B. NCD: no counter-defense, CD: counter-defense, BH: bet-hedging.

| Pathogen | Host |  |  |
| --- | --- | --- | --- |
|  | Phenotype 1 (ND) | Phenotype 2 (BH) | Phenotype 3 (D) |
| Phenotype 1 (NCD) | 9 | 1.86 | 0.67 |
| Phenotype 2 (BH) | 8 | 6.81 | 6.61 |
| Phenotype 3 (CD) | 7 | 6.35 | 6.24 |

Long-term fitness in a fluctuating environment is determined by the geometric mean (Simons, 2011). Bet-hedging strategies maximize geometric mean fitness across all generations (Haaland *et al*., 2019). We calculate the means and variances in phenotype that maximize arithmetic or geometric mean fitness in interaction with different phenotypes of pathogens. Now, we determine the fitness mean for the entire population (Table 6).

**Table 6.** Fitness matrix for host and pathogen population. AM: Arithmetic mean, GM: Geometric mean.

|  | Host |  | Pathogen |  |
| --- | --- | --- | --- | --- |
|  | AM | GM | AM | GM |
| Phenotype 1 (ND/NCD) | 3 | 3 | 3.84 | 2.24 |
| Phenotype 2 (Bet-hedging) | 4.99 | 4.26 | 7.14 | 7.11 |
| Phenotype 3 (D/CD) | 4.49 | 3.40 | 6.53 | 6.52 |

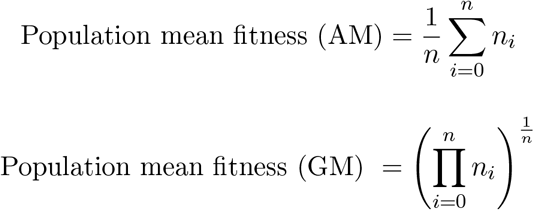

where *n* = number of phenotypes, AM = Arithmetic mean and GM = Geometric mean. Table 6 gives the mean fitness for different phenotypes of hosts and pathogens.

In Figure 10, an alternative bar plot is given for the mean fitness of the host and pathogen. For the illustrative parameter set, the intermediate phenotype 2 has higher arithmetic and geometric mean fitness across the considered interaction contexts (Table 6). As a result, we can say that the use of defense or counter-defense with a varying amount of toxin or enzyme as a bet-hedging strategy to increase fitness for the host and pathogen would be beneficial when interactions are uncertain (Figure 10).

**Figure 10.**
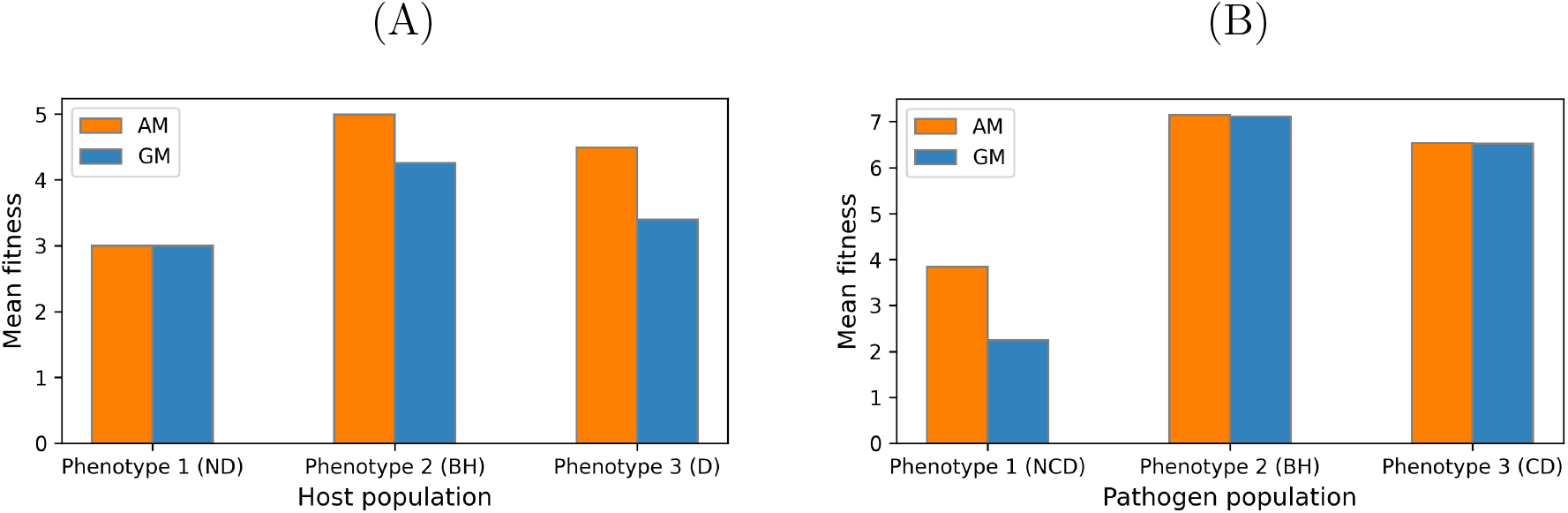
Mean fitness for all phenotypes of host and pathogen population. AM: Arithmetic mean (orange), GM: Geometric mean (blue)

## 3 Discussion

Here, we analyzed the dynamics in defense and counter-defense games between hosts and pathogens to generate models based on ordinary differential equations. In this asymmetric game, host and pathogen are players, where defense, counter-defense and no response are their strategies (Dwivedi *et al*., 2025). We built three models in this paper. In Model 1, we use replicator dynamics (Smith, 1982; Weibull, 1995; Hofbauer and Sigmund, 1998; Gintis, 2009; Cressman and Tao, 2014) to describe the interaction between a host and a pathogen and to analyze how an individual switches its strategies over time. We assumed that the host may produce a defense chemical (e.g. toxin) against the pathogen and the pathogen may produce a counter-defense chemical (e.g. enzyme) as a response to the host’s defense (Ewald *et al*., 2020; Dwivedi *et al*., 2025), where both actions require certain costs.

Our results explain continuous switching from no defense/counter-defense to defense/counter-defense and vise-versa for host and pathogen. Strategies oscillate permanently in the case of high benefit-to-cost ratio. In contrast, host and pathogen prefer no response for the low benefit-to-cost ratio, whether the interaction is between individuals or among populations (Dwivedi *et al*., 2025).

In Model 2, we analyze the game between host and pathogen populations, where a number of hosts and pathogens choose the defense and counter defense strategy, respectively, and they interact each other randomly. We formulated a model based on Lotka-Volterra equations (Hofbauer and Sigmund, 1998; Gintis, 2009; Ogata, 2010) to analyze how the host and pathogen populations change over time.

In Model 3, bet-hedging strategies for host and pathogen have been analysed. These resemble optimal mixed strategies in evolutionary game theory. Both bet-hedging and mixed strategies have the potential to lead to a diversity of phenotypes in the population (Li *et al*., 2017; Haccou and Iwasa, 1995). A bet-hedging strategy enables populations to survive in dynamic and uncertain habitats with a particular fitness cost (Morawska *et al*., 2022). While bet-hedging is often studied by stochastic approaches (Cohen *et al*., 1966; Seger and Brockmann, 1987; Starrfelt and Kokko, 2012), we have here used a deterministic approach (Cohen *et al*., 1966; Shemesh *et al*., 2013). We have determined the fitness of host and pathogen as a function of toxin and enzyme concentrations, respectively, based on Michaelis-Menten kinetics (Simms and Rausher, 1987; Siemens *et al*., 2010). This is compatible with an interpretation in terms of risk-spreading, but does not by itself constitute a full stochastic bet-hedging analysis.

The replicator and Lotka-Volterra equations in Models 1 and 2, respectively, are autonomous, nonlinear and deterministic in nature (Hofbauer and Sigmund, 2003). Replicator equations describe how individuals or populations switch their strategies over time based on their payoffs or fitness in a game (Weibull, 1995; Cressman and Tao, 2014). We obtained marginally stable periodic oscillations in Model 1 and irregular oscillations in Model 2. Since in the high-benefit case, the equilibrium points of these models are centers, all trajectories of the system are periodic orbits, and the periodic orbit can be evolutionarily stable in case of discrete replicator dynamics (Mukhopadhyay and Chakraborty, 2020). It is worth mentioning that the resulting marginally stable oscillations are (at least partly) due to the methods of replicator and Lotka-Volterra equations used here. Using other methods may lead to a stable steady state as, for example, when using game theory and considering three strategies (Dwivedi *et al*., 2023).

Varying time behavior in host-dynamics were also found by Buckingham and Ashby (Buckingham and Ashby, 2022; Kim and Ashby, 2026). Depending on the specificity between host and pathogen, stable monomorphism, polymorphism or oscillatory dynamics were found. An experimental evidence of coevolutionary dynamics within *Pseudomonas aeruginosa* PAO1 and a set of bacteriophages was found where four of six phages revealed, in time-shift assays, temporal peaks in bacterial resistance and and phage infectivity, pointing to fluctuating selection dynamics (Red Queen dynamics). For two phages, increased bacterial resistance ranges over the entire length of the experiment were observed (Betts *et al*., 2014).

The advantage of an autonomous system is that one can determine the equilibria and perform stability analysis more easily. Several methods (qualitative theories) exist to analyze the asymptotic stability of equilibria, such as Jacobian matrix, nullclines (Murray, 2002; Otto and Day, 2007) and Lyapunov stability (Sandholm *et al*., 2010; Hofbauer and Sandholm, 2009).

It is worth noting that mixed Nash equilibria and, thus, oscillations, can arise both in symmetric and asymmetric games. The number of strategies necessary is two and three, respectively. The matching pennies game (Tadelis, 2013) (with two strategies) and the rock-paper-scissors game (Gintis, 2009) represent instructive examples. The two-player rock-paper-scissors game can even lead to Hamiltonian chaos (Sato *et al*., 2002).

The game-theoretical framework for host–pathogen defense and counter-defense interactions is part of a broader program applying evolutionary game theory to biological conflicts across scales and systems. Analogous game-theory models in cancer show, on the one hand, phenotypic trade-offs formalized as a go-or-grow game (migration vs. proliferation) (Basanta *et al*., 2008; Dwivedi *et al*., 2023). On the other hand, it explains how costly diffusive signaling architectures can suppress exploitative mutants by lowering their fixation probability (Oña and Lachmann, 2020). These parallels suggest that the logic of defense and counter-defense, where costly investments yield frequency-dependent benefits, is a general feature of antagonistic coevolution, whether between species or within tissues.

Our model provides the scenario to show how the host and pathogen evolve their defense and counter-defense response as their bet-hedging strategies. In the present model, the arithmetic mean and geometric mean fitness of the bet-hedger population are higher than that of the non-bet-hedger population. A disadvantage is that to maximize the geometric mean, an individual has to reduce its fitness variance (Olofsson *et al*., 2009). For example, a summer specialist has a higher fitness in summers compared to winters, while a generalist has about the same fitness contributions in both seasons (Olofsson *et al*., 2009). For simplicity, we ignored fitness variance in our model and considered the efficacy of producing defense or counter-defense as an individual fitness. Another example is the ‘adaptive coin flipping’ strategy in the decision-theoretical examination of natural selection, where a third strategy can evolve under a variety of conditions (Cooper and Kaplan, 1982) as a bet-hedging strategy.

In conclusion, we can say that there is always a switch between strategies with a certain amount of defense (toxin) or counter-defense (enzyme). The famous “Red Queen hypothesis” in evolutionary biology asserts that organisms consistently adapt and evolve to survive against ever-evolving pathogenic organisms (Buckingham and Ashby, 2022). A related example is thoroughly explained in plant-pathogen interactions (Clay and Kover, 1996). A special case of such a dynamics is represented by red-queen cycles (Buckingham and Ashby, 2022; Kim and Ashby, 2026). In the future, it will be useful to refine the present model using further modeling methods such as stochastic approaches and to verify the results with experimental data.

## Acknowledgements

Stimulating discussions with Albert A. Pinto (Porto, Portugal), Rosalind Allen and Lukas Korn (Jena) are gratefully acknowledged. S.D. thanks the Jena School of Microbial Communication (JSMC) for support. The authors used Google Gemini (version 3) for text optimization. The authors reviewed and edited the content as needed and take full responsibility for the content of the article.

## Author Contributions

S.D. conceptualized the study, established the mathematical model, performed the simulations, produced the figures and wrote the manuscript. S.S. supervised the study and refined the model. L.O. conceptualized part of the the study and performed simulations. All authors verified the results and reviewed the manuscript.

## Availability of Data and Materials

All data generated or analyzed during this study are included in this article.

